# Neuropeptide relay between SIFa signaling controls the experience-dependent mating duration of male *Drosophila*

**DOI:** 10.1101/819045

**Authors:** Kyle Wong, Justine Schweizer, Khoi-Nguyen Ha Nguyen, Shatha Atieh, Woo Jae Kim

## Abstract

*Drosophila melanogaster* is a suitable model for investigating how neuropeptides influence animal behaviours and physiology. We previously reported that two behavioural paradigms control mating duration of male *Drosophila*, called Longer-Mating-Duration (LMD) and Shorter-Mating-Duration (SMD) that are induced through socio-sexual environment prior to copulation. Understanding the molecular and cellular mechanisms by which males exhibit plasticity to different social cues remains poorly understood. Here, we show that SIFa modulates the neural circuitry for both LMD and SMD. Neuropeptide-to-neuropeptide communication, so called ‘neuropeptide relay’ plays a key role to mediate this control. We identified that 7 neuropeptides expressed in SIFa Receptor-positive cells are functionally important to regulate either LMD and/or SMD. The modulation of two independent mating duration behaviour by the different SIFa-mediated neuropeptide relay will help to further investigate how the neuropeptidergic modulation can control complex behaviours.

## INTRODUCTION

Neuropeptides represents a diverse class of signaling molecules that modifies neural activities and neural circuit dynamics. One of its fundamental features is its ability to modulate the properties of neural circuits (i.e. decision-making circuits and/or pattern generators) by changing neuronal excitability causing them to be multifunctional [1]. Neuromodulators are coupled into highly complex networks, from which distinct behavioural qualities emerge as a basis of these functions [2].

Neuropeptides constitutes a large and diverse class of signaling molecules that are produced by various types of neurons, which modulates many insect behaviours and physiology [3,4]. Recent advances in neurobiological research have suggested that neural circuits encompassing neuropeptide relay signaling are essential modulators for various behaviours, including feeding, energy homeostasis, sleep and wakefulness [5–11]. It has been well know that appetite and satiety work together to regulate the balance of animal’s physiology and behaviour through neuropeptide relay. For example, the hypothalamus in mouse brain functions as a central regulator in satiety control using two opposing types of neurons, orexigenic and anorexigenic neurons of which are distinctively characterized by the expression of specific neuropeptides such as NPY/AgRP or POMC/CART [6].

It has been recently reported that male-specific mating behaviours are also modulated by neuropeptide relay [13–15]. In *Drosophila*, males respond to potential threats in mating competition by altering their behaviour prior to copulation. One of these strategies is the male’s ability to accurately control mating duration. We reported that the rival-induced extended mating, Longer-Mating-Duration (LMD) [13,14] and sexual experience-based shortened mating, Shorter-Mating-Duration (SMD) of *Drosophila* provide powerful models for investigating neuropeptidergic systems as its molecular and cellular mechanisms. LMD signaling relies on two neuropeptides, PDF and NPF, which is modified by the male’s prior experience with other rival male flies [13,14]. In contrast, SMD signaling is modulated by prior sexual experience which induces neuronal changes to sNPF neurons [15]. Despite much evidence indicating that neuropeptidergic systems are actively involved, how these modulatory effects result in circuit-level consequences is still not well understood.

The neuropeptide AYRKPPFNGSIFamide or SIFa, is a neuromodulator that demonstrates plasticity on a molecular and behavioural level. SIFa was first identified in grey fleshfly *Neobellieria bullata* [16–18] and is expressed by four neurons located in the pars intercerebralis which is well distributed throughout the brain and VNC by its morphology [19]. This neuropeptide has significant impact on various behaviours including appetite [12], courtship [17], and sleep [20]. However, the mechanism of SIFa signaling controlling various behaviours is not well understood.

Here, we report the relationship between the SIFa-mediated signaling and its functional relevance to LMD and SMD in *D. melanogaster*. The modulation of two independent behavioural paradigm controlling mating duration by SIFa-mediated neuropeptide relay will help to further investigate how the neuropeptidergic modulation can control complex behaviours.

## METHODS

### Drosophila Stock Lines

All flies were raised in 12:12 light to night conditions for all experiments. Each fly is collected between 0-3 days after eclosion are assigned to either the LMD or SMD paradigm. Once collected, male flies are retained in same environment until further manipulations are needed. LMD conditions contain group-reared (naïve) and singly-reared flies whereas SMD conditions contain group-reared (naïve) and sexually experienced-reared.

### Mating Duration Assay / Mating Assay

The mating duration assay in this manuscript has already reported elsewhere [13–15]. In this manuscript, we merged LMD and SMD assay together. In short, to setup mating duration assay, measurement for LMD is calculated between group-reared (naïve) and singly-reared flies whereas measurement for SMD is calculated between group-reared (naïve) and sexually experienced-reared. The relative significant difference between each groups determines whether each behaviour is exhibited or not. To setup experiment, each fly is put to sleep using CO2 system then assigned to mating chamber which contains up to 36 pairs of flies. When performing assay, only flies that mate for at least 10 minutes are included in total results.

### Immunohistochemistry

The dissection and immunostaining protocols for the experiments are described elsewhere [13–15]. In short, the *Drosophila* brain will be extracted from adult flies after 5 days of eclosion and fixed into 4% formaldehyde for 30 minutes at room temperature. The sample will be washed in 1% PBT three times (30 minutes each) and blocked in 5% normal donkey serum for 30 minutes. The sample will then be incubated with primary antibodies in 1% PBT at 4°C overnight followed by the addition of fluorophore-conjugated secondary antibodies for 1 hour at room temperature. The brain will be placed onto an antifade mounting solution (Invitrogen) on slides for imaging.

### Statistical Tests

All statistical tests were conducted using Graphpad Prism 5. Analysis of variance (ANOVA) was conducted for all experimental conditions and Tukey’s test to determine significance within groups examined. For internal control, we performed experiments using wild-type for each experiment for internal comparison. (*** = p < 0.001, ** = p < 0.01, * = p < 0.05).

## RESULTS

### SIFa acts as a ‘molecular switch’ between LMD and SMD

To investigate how SIFa signaling modulates LMD and SMD of male fruit flies, we merged the two behavioural paradigms of LMD and SMD as we previously described [13–15]. Thus, male^*naive*^ represents group-reared condition, male^*single*^ represents singly-reared, and male^*exp.*^ represents group-reared males with prior sexual experience (The first panel in Fig. 1A). The difference between male^*naive*^ and male^*single*^ represents LMD behaviour and difference between male^*naive*^ and male^*exp.*^ SMD behaviour. When we found the differences in statistical significant manner, we explain it as LMD and/or SMD behaviour is ‘intact’. However when we found the difference between groups is non-significant, we explains it as LMD and/or SMD behaviour has ‘disrupted’ or ‘disappeared’ [13–15].

**Figure 1.**
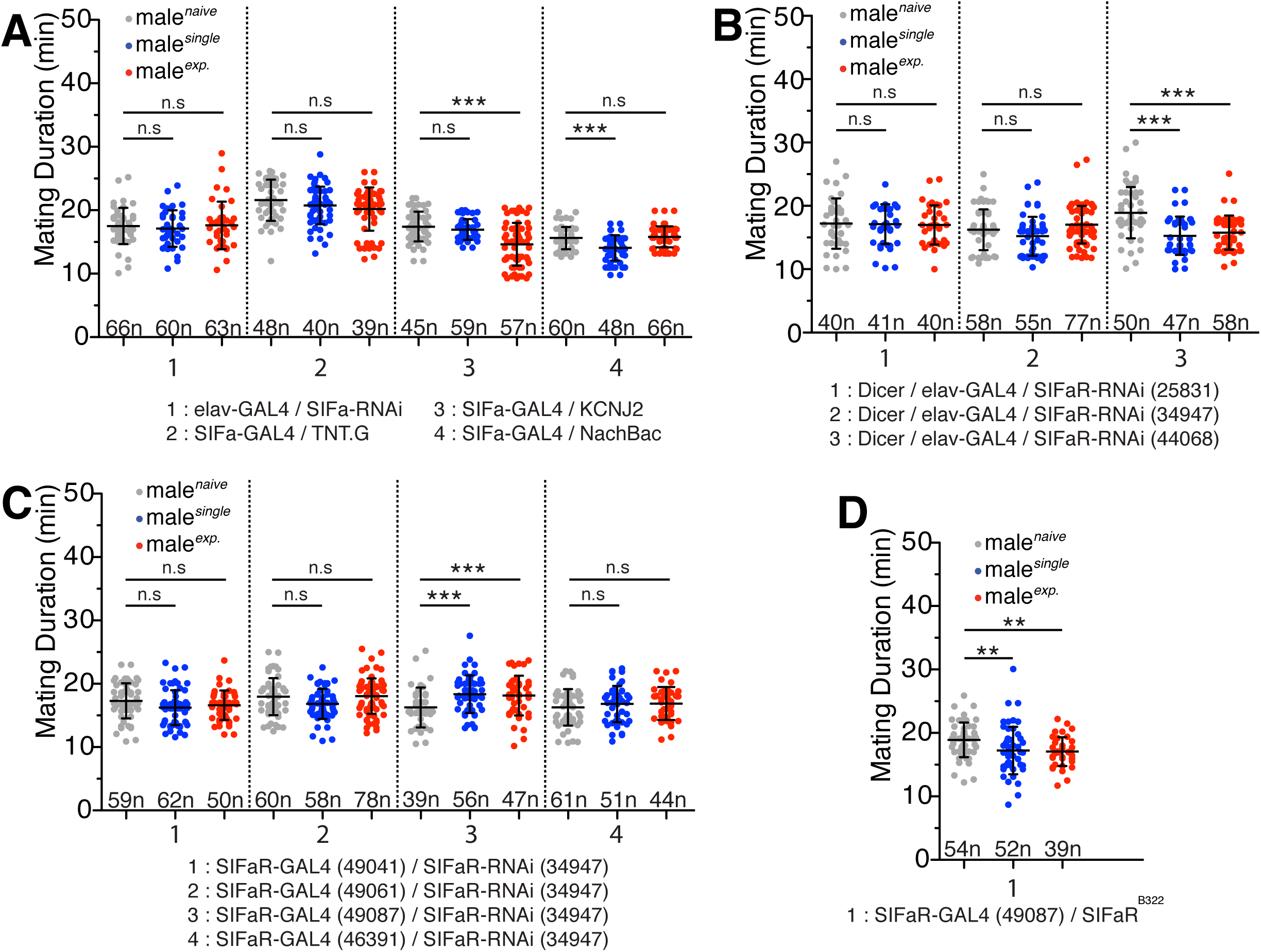
SIFamide signaling is required for both LMD and SMD behaviour. (A) SIFa modulates the activation between both LMD and SMD male behaviour. Downregulation of SIFa in elav-positive cells displayed a disruption in both LMD and SMD. Insertion of synaptic blocker, TNT, in SIFa neurons disrupted both LMD and SMD. Preventing membrane depolarization via KCNJ2 disrupted LMD but not SMD. Increasing sodium conductance via NaChBac disrupted SMD but not LMD. (B) Distinct SIFaR-RNAi expression in neuronal-specific tissue disrupts both LMD and SMD. SIFaR-RNAi 25831 and 34947 but not 44068 effector lines disrupted both LMD and SMD. (C) Knockdown using SIFaR-RNAi in different subsets of SIFaR-positve cells disrupts both LMD and SMD. Downregulation of SIFaR in SIFaR-GAL4 49041, 49061, 49087, and 46391 driver lines disrupted LMD and SMD. (D) LMD and SMD was rescued using SIFaR-GAL4 49087 in SIFaRB322 mutant phenotype. (A-D) Each dot in the box and whisker plot (n) signifies a single pair (Male/Female) of flies. Statistical analysis was conducted using one-way ANOVA followed by Kruskal-Wallis comparisons test. Numbers at x-axis indicates n. NS indicates non-significance (p>0.05), ** indicates significance (p<0.01) and *** indicates significance (p< 0.001) for comparisons to naive group. Error bars indicate SD.

To understand the function of SIFa in LMD and SMD, we knocked down SIFa neuropeptide using SIFa-RNAi with pan-neuronal *elav-GAL4* driver. The results showed that pan-neuronal knockdown of SIFa disrupts both LMD and SMD (The first panel in Fig. 1A). Neuropeptides are generally packaged in large dense-core vesicles, and co-existed with neurotransmitters in small synaptic vesicles [21]. To confirm whether LMD and/or SMD is controlled by synaptic vesicle transport, we expressed tetanus toxin light chain (TNT) that cleaves synaptobrevin (nSyb) and block the synaptic transmission [22]. The expression of *UAS*-TNT by SIFa-*GAL4* results a loss of both LMD and SMD, suggesting that the synaptic vesicle transport is required to drive both LMD and SMD (The second panel in Fig. 1A).

Next, we manipulated the electrophysiological functions of SIFa-positive neurons by expressing potassium channel KCNJ2 to constitutively hyperpolarize target neurons and found that only LMD but not SMD was disrupted (The third panel in Fig. 1A). In contrast, when we constitutively depolarized the same neurons by expressing bacterial sodium channel NachBac-GFP, we found that only SMD but not LMD was disrupted (The last panel in Fig. 1A). Our findings indicate that SIFa neurons acts as a ‘molecular switch’ between LMD and SMD since the hyperpolarization of SIFa disrupts LMD whereas the depolarization disrupts SMD.

To confirm whether LMD and/or SMD are dependent on SIFa-mediated signaling through SIFamide receptor (SIFaR), we used three different SIFaR-RNAi lines and determined the efficacy of each line. We combined each RNAi line with pan-neuronal elav-*GAL4* driver and found a loss of both LMD and SMD in two of three SIFaR lines (Fig. 1B). Of those two lines, we decided to use SIFaR-RNAi 34947 for future knockdown experiments.

To determine which subpopulations of SIFaR-positive cells are sufficient for LMD and/or SMD, we selected four available SIFaR-*GAL4* lines. Since each of the selected lines utilized different fragments of the genomic DNA of the SIFaR promoter, it allows us to target different subpopulations of SIFaR-positive cells [23]. Knockdown of SIFaR with all SIFaR-*GAL4* drivers can disrupt both LMD and SMD (Fig. 1C), suggesting that the expression of SIFaR in SIFaR-*GAL4* labeled cells are necessary for both LMD and SMD behaviours. Interestingly, knockdown of SIFaR with SIFaR-*GAL4* (49087) driver results longer mating duration in singly reared and in experienced condition than naïve one (The third panel in Fig. 1C).

To determine which SIFaR-positive cells are sufficient for LMD and SMD, we designed genetic rescue experiments using a mutant SIFaR (*SIFaR*^*B322*^) strain that are homozygous lethal. To rescue the lethality and behaviour, we attempted to restore LMD and SMD by reintroducing the wild type SIFaR gene into SIFaR mutant background using SIFaR-*GAL4s*. The results showed that two of four SIFaR-*GAL4* lines produced homozygous adults, suggesting that those two *GAL4* expressing cells (49061 and 49087) are essential for the lethality during development (Table 1). We subsequently used both lines for mating duration assays and found that only SIFaR-*GAL4* 49087 driver can rescue both LMD and SMD behaviour (Fig. 1D). These findings suggest that SIFaR-*GAL4* driver 49087 labels the cell populations that are necessary and sufficient for both LMD and SMD behaviours via SIFa-SIFaR signaling pathway.

The bHLH transcription factor DIMMED has been associated with the differentiation of peptidergic cells in *Drosophila* [24] and the most of the neuropeptidergic cells are labeled by DIMM-*GAL4*. To confirm that SIFaR-positive cells are also neuropeptidergic cells that are expressing specific neuropeptides, we knocked down SIFaR using DIMM-*GAL4* strain and found that only SMD disappears (Fig. 2A). This data suggests that SIFaR signaling in DIMM-positive cells are only required for SMD but not LMD behaviour.

**Figure 2.**
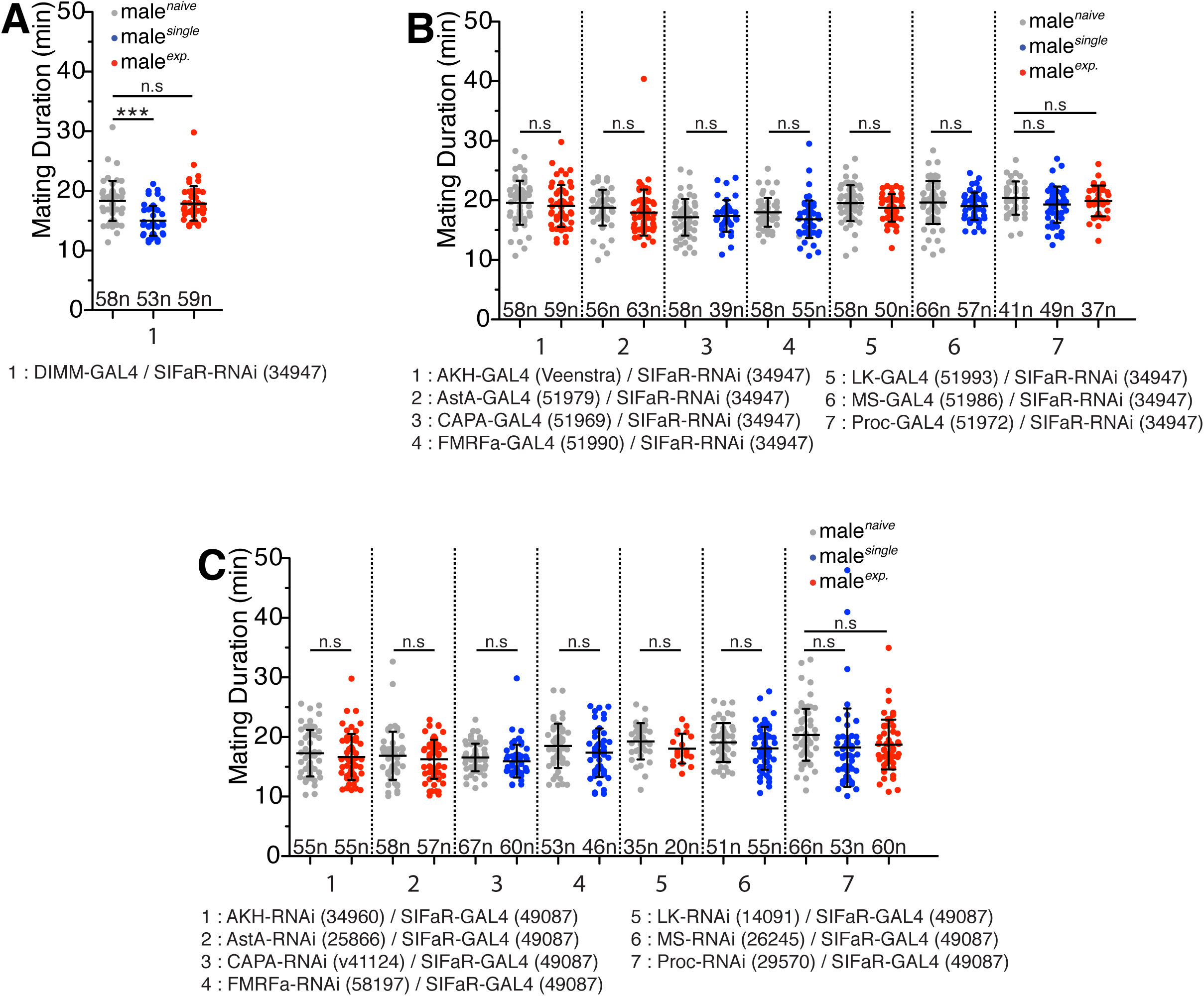
Identification of candidate neuropeptidergic neurons that regulate either LMD and/or SMD. (A) Knockdown of SIFaR in C929-*GAL4* (DIMM) labeled cells. (B) Knockdown of SIFaR in different neuropeptide-*GAL4* labeled cells. (C) Knockdown of candidate neuropeptides in SIFaR-*GAL4* labeled cells. (A-C) Each dot in the box and whisker plot (n) signifies a single pair (Male/Female) of flies. Statistical analysis was conducted using one-way ANOVA followed by Kruskal-Wallis comparisons test. Numbers at x-axis indicates n. NS indicates non-significance (p>0.05), ** indicates significance (p<0.01) and *** indicates significance (p< 0.001) for comparisons to naive group. Error bars indicate SD.

Systematic *GAL4* screening is a powerful method for investigating the relationship between the subpopulation of cells and complex behaviours [25]. To identify potential candidate neuropeptidergic neurons that express SIFaR which are required for LMD and/or SMD, we conducted a neuropeptide-*GAL4* screening using SIFaR-RNAi. We analyzed total 19 different neuropeptide-*GAL4* strains [26] and summarized the results in Table 2. In this screen, we found that knockdown of SIFaR in proctolin-positive neurons disrupted both LMD and SMD, knockdown of SIFaR in CAPA-, FMRFa-, and MS-positive neurons disrupts only LMD not SMD, and knockdown of SIFaR in AKH-, AstA-, and LK-positive neurons disrupts only SMD not LMD (Fig. 2B). These data suggest that three neuropeptides (CAPA, FMRFa, and MS) may be only required for LMD which are DIMM-negative and three neuropeptides (AKH, AstA, and LK) may be only required for SMD which are DIMM-positive.

To confirm whether the candidate neuropeptides are functionally important to regulate either LMD and/or SMD, we knocked down candidate neuropeptide with proper RNAi in relevant SIFaR-positive cells. In this screen, we used a total of 28 different candidate neuropeptide-RNAi lines with SIFaR-*GAL4* 49087 driver and examined for changes in both LMD and SMD (Table 2). Our results show that seven neuropeptides identified through *GAL4* screen are functionally required for LMD and/or SMD behaviour (Fig. 2C). In summary, we functionally confirmed that proctolin requires for both LMD and SMD behaviours, CAPA, FMRFa, and MS requires for LMD but not SMD, and AKH, AstA, and LK requires for SMD but not LMD.

To visualize the candidate neurons expressing various neuropeptide and SIFaR together, we used two independent binary systems, *GAL4*/*UAS* and LexA/LexAop [27]. Since neuropeptide screening was conducted via *GAL*/*UAS*, we visualized SIFaR cells via *LexA*/*LexAop* system. First, we performed experiments to confirm whether SIFaR-*LexA* expression pattern is same or similar to SIFaR-*GAL4* expression pattern. We expressed mCD8GFP via SIFaR-*LexA* and mCD8RFP via SIFaR-*GAL4* and revealed similar morphology in central brain and few variations in the VNC region (Fig. 3A). This finding demonstrates that the subpopulations of SIFaR cells via LexA is suitable to recapitulate the results from previous SIFaR-*GAL4* experiments.

**Figure 3.**
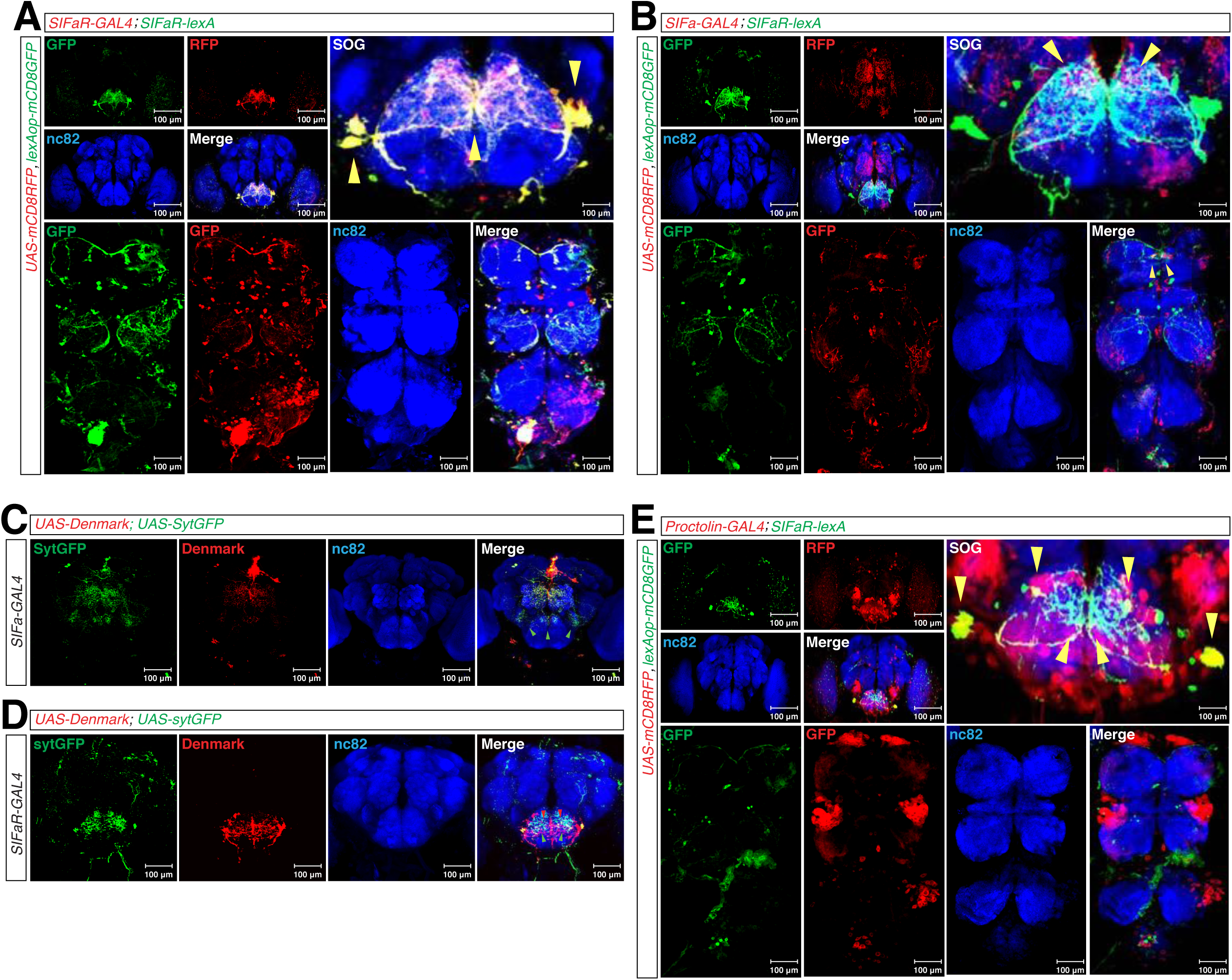
Two Pairs of SIFaR-positive and neuropeptidergic-positive neurons localized to the SOG region are critical to generate LMD and SMD. (A) Expression pattern of a SIFaR-*LexA* labeled cells (green) and SIFaR-*GAL4* labeled cells (red) using membrane tag marker (yellow arrowheads indicate overlap). (B) Expression pattern of a SIFaR-*LexA* labeled cells (green) and SIFa-*GAL4* labeled cells (red) using membrane tag marker (yellow arrowheads indicate potential synapses). (C) SIFa-*GAL4* labeled cells dendritic and axon terminal expression pattern (green arrowheads indicate axon terminals). (D) SIFaR-*GAL4* labeled cells dendritic and axon terminal expression pattern (green arrowheads indicate axon terminals and red arrowheads indicate dendrites). (E) Expression pattern of SIFaR-*LexA* labeled cells (green) and proctolin-*GAL4* labeled cells (red) using membrane tag marker (Yellow arrowheads indicates potential synapses).

Next, we visualized the expression pattern of SIFa-SIFaR signaling pathway to find out their synaptic contacts. To do this, we labeled SIFaR-positive cells using the membrane marker mCD8GFP via SIFaR-lexA and SIFa neurons using the mCD8RFP via SIFa-*GAL4*. Our findings highlight extensive SIFa and SIFaR arborization in the medial central brain and VNC (Fig. 3B). Interestingly, we identified an overlapping pattern restricted to the subesophageal ganglion (SOG) region in the brain and to the superior region in the VNC which indicates potential synapses (yellow arrows in Fig. 3B). This data indicates that SIFa neurons form multiple synapses with SIFaR neurons in the brain and VNC.

To provide more evidence that the overlapping region by SIFa-*GAL4* and SIFaR-*lexA* are genuine synapses, we expressed dendritic marker DenMark and presynaptic marker sytGFP via SIFa-*GAL4* (Fig. 3C) or SIFaR-*GAL4* (Fig. 3D). This data showed that SIFa-positive neurons have extensive dendritic and axon terminals throughout the entire central brain including the axon terminals in the lateral SOG region (green arrows in the Fig. 3C). In contrast, both of the dendritic arborizations and axon terminals of SIFaR-positive cells concentrated near SOG, especially in the central regions (green and red arrows in the Fig. 3D). We propose that a principle signaling pathway that mediates LMD and SMD is likely modulated by synapses formed with SIFa neurons with SIFaR-positive cells in central SOG region.

Next, we visualized the morphological distribution of each candidate neuropeptide-positive neurons by expressing mCD8GFP via SIFaR-*LexA* and mCD8RFP via seven candidate neuropeptide-*GAL4s*. For AstA, there is restricted bilalteral expression in central brain with DIMM expression in inferior VNC region (red arrows in the Fig. S1A). We found that no expression in AKH neurons in either central brain or VNC (Fig. S1B). Targeting CAPA neurons revealed expression in central brain including the optic lobes, SOG, and VNC regions (Fig. S1C). DIMM-positive CAPA neurons appear to be localized towards the medial superior VNC region (red arrows in the Fig. S1C). Expression of FMRFa neurons are restricted towards the optic lobe, SOG regions, and DIMM expression in VNC region (Fig. S1D). We found that FMRFa-positive cells overlaps with SIFaR-positive cells in optice lobe and subsequent SOG regions (yellow arrows Fig. S1D). For LK, we found expression in medial SOG region (Fig S1E). Our immunostaining data showed one cell expresses both SIFaR and LK in the medial SOG region (yellow arrow in Fig. S1E). For MS, we found some expression in the superior lateral region near the optic lobe (Fig S1F) and found small number of cells that express both MS and SIFaR in the optic lobe (yellow arrow in Fig. S1F). Finally, we found that proctolin-positive cells strongly overlap with SIFaR-positive cells in the central and lateral SOG region (yellow arrows in Fig. 3E), suggesting that proctolin is critical mediator of SIFa signaling that modulates both LMD and SMD behaviours.

By visualizing neurons that express both SIFaR and candidate neuropeptides, we showed that neuropeptide relay would be a critical way to modulate both LMD and SMD behaviours. However, these data might represent the limitation of our genetic tools to capture all the SIFaR-positive cells using *lexA* system (non-overlapping regions in the Fig. 3A) since we could not capture the overlapping neurons in the visualization process (Fig. S1). Constructing better SIFaR-lexA strains that can cover the most of SIFaR-positive cells would help to identify the further overlapping cells that express SIFaR and each candidate neuropeptide.

## Discussion

In this study, we examined the association of SIFa-mediated signaling and its functional relevance to male-specific mating behaviours, LMD and SMD in *D. melanogaster*. The experiments reported here identify two pairs of bilateral SIFa neurons in the pars intercerebralis of the adult brain are critical to switch between LMD or SMD behaviours (Fig. 1). We also found that SIFaR expression in SIFaR-*GAL4s* labled cells are critical for both LMD and SMD behaviours (Fig. 1). Intriguingly, SIFaR cells which are DIMM-positive are essential for LMD whereas SIFaR cells which are DIMM-negative are important for SMD (Fig. 2). We also performed neuropeptide *GAL4*-based screen to identify the SIFaR-expressing neuropeptidergic neurons which can mediate neuropeptide relay for each behaviour (Fig. 2 and Table 2). We found seven neuropeptide-positive neurons as well as neuropeptide themselves are involved in LMD and SMD behaviours through RNAi screens (Fig. 2 and Table 2). Finally, we conducted an anatomical-based screen to identify which candidate neuropeptides are co-labelled with SIFaR and found three pairs of bilaterally proctolin neurons in the SOG, which indicate that these cells receive specific neuropeptidegic input from SIFa that modulate both LMD and SMD and send output signals to further modulate LMD and SMD behaviours (Fig. 3). We also identified SIFaR-positive neuropeptide-expressing cells in FMRFa, LK, and MS (Fig. S1) We propose that SIFa-mediated signaling through different neuropeptidergic cells can provide an excellent model to describe the molecular mechanisms that how the neuropeptide relay modulate complex behaviours.

Previous studies have reported a link between SIFa and interval timing behaviours [12,28], but no prior studies have identified a direct connection to mating duration. Our study showed that depolarizing SIFa neurons initiates LMD while hyperpolarizing displays SMD. This is the first evidence providing a causal link between LMD and SMD whereby SIFa neurons modifies its intrinsic cellular properties in accordance to extrinsic factors such as socio-sexual conditions. We also report that knockdown or eliminating synaptic vesicle exocytosis with TNT blocker [22] in SIFa neurons disrupted both LMD and SMD which suggests that the neuropeptide itself is also essential. Moreover, we showed that neuropeptide relay via SIFa-SIFaR signaling is specifically involved in switching LMD and SMD behaviours.

Neuropeptide relay are fundamental for survival and goal-oriented behaviours. One apparent case is the balance between appetite and satiety via orexigenic and anorexigenic neuronal circuits. Appetitive signals activate specific sensory neurons that then turn on a cascade of neural connections that initiate and maintain stereotyped behaviours such as feeding. Persistent feeding behaviour induced by appetite then induce production of satiety signals that, in turn activates a cascade of neural circuits to induce stereotyped opposing behaviours such as eating cessation. The nodes of those neural circuits activated either by appetite or satiety signals presumably inhibit each other at some level to facilitate sharply delineated behavioural outcomes. It has been known that neuropeptides and hormones are important neural substrates for modulating appetite and satiety control [6,29]. SIFa has been known as a central modulator in parallel inhibition between orexigenic and anorexigenic pathways. By acute activation in SIFa neurons, non-starved flies act starve-like and enhance their food intake. Interestingly, SIFa integrates orexigenic signals from myoinhibitory peptide neurons and anorexigenic signals from Hugin-pyrokinin neurons to mediate satiety and hunger [12]. This report suggests that SIFa neurons are important to recruiting compensatory pathways via other neuropeptides to facilitate hunger-driven behaviour.

In another study, SIFa-mediated signaling facilitates sleep functions in the pars intercerebralis [20], possibly through diuretic hormone 44 neurons [30]. Therefore, SIFa neurons does not act independently, but rather coordinates its activity collectively in response to other neuropeptides dependent of the behaviours it modulates. We believe that this integration of neuropeptidergic signaling towards SIFa may also account for the time delay in our mating paradigms. LMD and SMD utilizes PDF, NPF, and sNPF signaling which requires long but lasting effects to activate their corresponding receptors [31], this coincides with the one day delay to elicit either LMD or SMD [13–15]. To elucidate how this mechanism might occur, further studies will be required to understand the interactions between PDF/NPF and sNPF to SIFa neurons.

Here, we found that knockdown of SIFaR through SIFaR-*GAL4* (49087) driver-labeled cells display increased mating duration in singly reared and in experienced reared than naïve condition, thus representing a possible neuronal disinhibition circuit. In *Drosophila*, behavioural disinhibition has been observed in the mushroom body (MB) αβ and γ neurons [32] via MB247-*GAL4* [33]. Our previous work revealed that neuronal activity from OK107-*GAL4* driver-labeled MB cells are required for SMD. OK107-*GAL4* neurons appear to overlap with αβ and γ neurons [34], hence may play a role in our underlying disinhibition circuit. MB cells expressed by OK107-*GAL4* driver are cholinergic [35] and converge to MB output neurons (MBONs) which have afferent dopaminergic neurons (DANs) [36,37]. At OK107-MBONs synapses, sNPF potentiates Ach-evoked response in MBONs which has significant effects for odor-associated memory formation [35]. This is important for our paradigm as SMD is highly dependent on previous mating or prior learning experiences. Modulation of this circuit might further be adjusted by DANs. Interfering with dopaminergic signaling induces disinhibition [32,33], thus it is conceivable that target neurons may also express SIFaR. When constitutively activating these central neurons, this mediates pleiotropic effects or disinhibition as seen in our observations.

Our findings also report that bHLH transcription factor DIMM is essential to modulate SMD but not LMD. We also show that all candidate neuropeptides to be DIMM-positive and distributed throughout the entire nervous system. DIMM is expressed by large cells that show episodic release of amidated peptides (LEAPs) [24]. It remains to be seen how SMD might be modulated by one or all DIMM-positive neuropeptides, but we propose that insulin producing cells (IPCs) interact with AKH, AstA, and LK to mediate SMD (Fig 4.)

**Fig 4.**
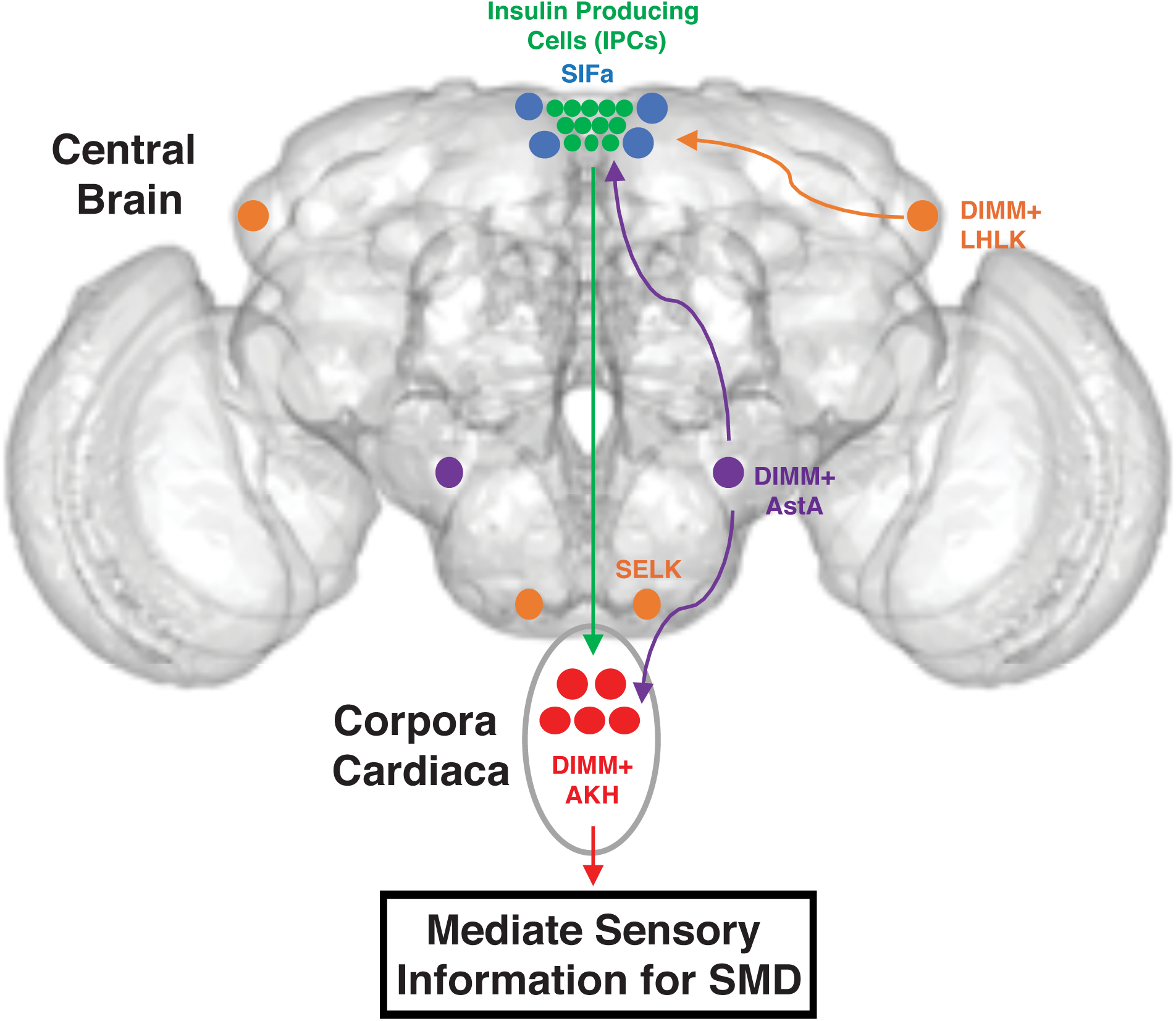
Proposed pathway where AKH, AstA, and LK modulates SMD through DIMM. Our proposition includes that SMD behaviour is driven by a current hunger-driven circuit that becomes incorporated into mating duration-specific behaviour. The mechanism is unknown but all of our neuropeptides that modulates SMD are potentially DIMM-positive.

Previous studies have described two types of LK neurons, lateral horn leucokinin (LHLK) and subesophageal ganglion leucokinin (SELK) neurons. LHLKs are linked with locomotor activity, metabolism, and sleep [38] but its distinct role is still unspecified. In one study, an increased in growth was observed in DIMM-positive LHLK neurons which are controlled by *Drosophila* insulin receptor via insulin/IGF signaling in adult flies [39] and interact with IPCs [40]. Our results selectively targeted LHLK neurons with our DIMM and LK knockdown result on SMD, thereby may indicate a novel function in LHLK-IPC interactions (Fig. 2 and 3). Conversely, our expression data indicates that SELKs modulate SMD via SIFaR and may relay the signal towards the IPC (Fig. S1). There are no reports of synaptic contact between these IPCs and SELKS [41], suggesting that LK influences IPCs in a paracrine manner. However, the IPCs are in synaptic contact with LHLKs [41] and are also modulated by octopamine [42], serotonin [40], and sNPF where its dendrites branch to the pars intercerebralis and tritocerebrum and axonal terminals in the corpora cardiac and anterior aorta and midgut region [43]. Interestingly, we previously found one sNPF neuron localized as dorsolateral peptidergic neurons (DLPs) to be robustly active in experience males [15], these DLPs are also thought to regulate IPC activity [43]. Though mechanistic evidence is lacking, we propose that IPCs interacts with both LHLKs and/or SELKs and contributes to integrating modulatory effects from other neuropeptidergic pathways to elicit SMD.

SMD behaviour may also be mediated by AKH and AstA acting on IPC circuits. AKH regulates energy storage homeostasis by mobilizing glucose and lipids while *Drosophila* insulin-like peptide (DILPs) is involved glucose metabolism [44]. When functioning together, they modulate glucose sensitivity for various behaviours including feeding [45], foraging [46], and sexual attractiveness [47]. Insulin signaling acts through AKH Receptors (AKHRs) which are expressed are also found in Gr5a gustatory neurons [44]. Our findings report that male flies deficient for AKH in SIFaR-GAL4 labelled cells had disrupted SMD behaviour. As AKH co-activates with insulin-like peptides to regulate glucose and lipid metabolism [48], we infer that AKH adjusts the dynamic balance of glucose levels in sensory Gr5a neurons. In fruit flies, memory can be reinforced up to several days when specific odors are paired with glucose [49]. We believe that the memory system required for SMD is interrupted and the male flies are no longer primed for this behaviour. This process is also facilitated through AstA-AKH interactions to maintain the dynamic balance of glucose. AstA is a neuropeptide involved in a series of satiety signals such as feeding, metabolism, and sugar reward [50–52]. Its receptor, Allatostatn A Receptor (AstAR), is expressed in both insulin and AKH cells. When silencing AstAR cells, AKH signaling was reduced which thus accumulated high lipid levels [51]. The authors claim that AstA is involved in nutrient homeostasis by controlling metabolism and feeding decisions [51]. In our case, downregulation of AstA in SIFaR-GAL4 labelled cells disrupted the metabolic balance that maintains glucose ratios through both IPCs and AKH signaling. This subsequently interrupts sensory integration to the appropriate systems and thereby inhibits SMD behaviour.

We propose that SMD signaling recruits neuronal circuits within this network via AKH, AstA, and LK by SIFaR to elicit this behaviour. In contrast, LMD appears to be solely independent of this pathway where it employs specialized microcircuits through CAPA and FMRFa to display this behaviour. In summary, our findings highlight a potential novel signaling mechanism where SIFa-mediated neuropeptide relay is required to modulate both LMD and SMD. Our study will help to investigate the mechanisms how does neuropeptide relay modulate specific behaviours.

---

**Table 1. Rescue mutant *SIFaR***^***B322***^ **behaviour by using different *SIFaR GAL4* lines.** Wild type SIFaR was reintroduced into *SIFaR*^*B322*^ mutant background using different SIFaR-GAL4 drivers (Bloomington stock number #49041, #49061, #49087, and #46931). Genotypes for each cross are as below; CS; UAS-SIFaR/CyO; *SIFaR*^*B322*^/TM6B X CS;; SIFaR-GAL4s, *SIFaR*^*B322*^/TM6B (third chromosome recombined strains).

**Table 2. Screening results of neuropeptide GAL4 and RNAi for SIFa signaling.** The first screens used SIFaR-RNAi (34947) crossed with different neuropeptide-GAL4 drivers. The second screens used SIFaR-GAL4 (49087) crossed with neuropeptide-RNAi strains. The results show that LMD and/or SMD are intact (marked with *, **, ***) or disrupted (n.s). No data suggests that the crosses were lethal or males could not succeed copulation thus mating duration assay could not be completed.

**Supplementary Fig 1.**
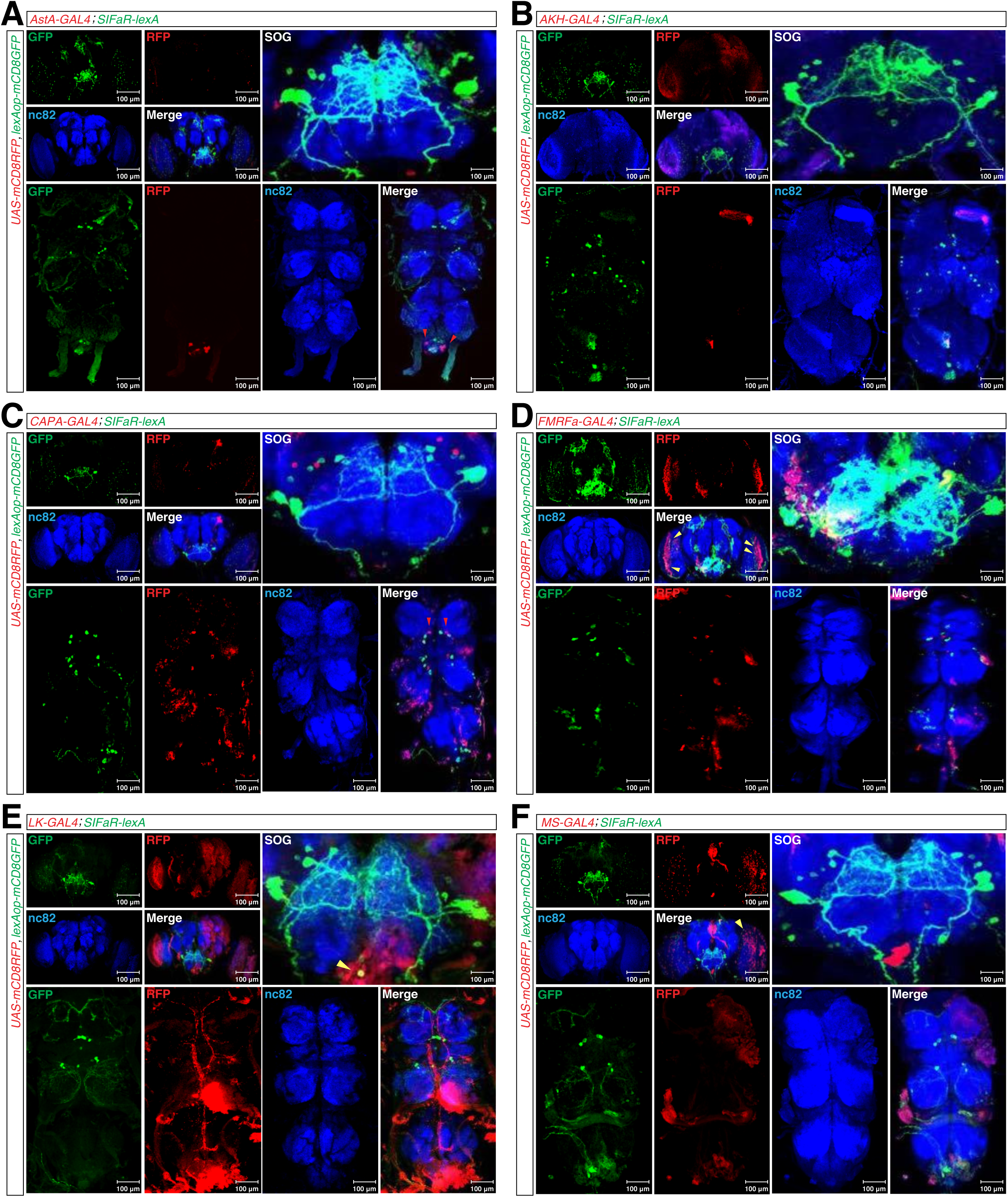
(A-F) Expression pattern SIFaR-LexA labeled cells (green) and candidate neuropeptide-GAL4 neurons (red) using membrane tag marker. (A) AstA (red arrowheads indicate DIMM expression). (B) AKH. (C) CAPA (red arrowheads indicate DIMM expression). (D) FMRFa (yellow arrowheads indicate overlap). (E) LK (yellow arrowheads indicate overlap). (F) MS (yellow arrowheads indicate overlap).

