## Supplementary material for "Neuropeptide relay between SIFa signaling controls the experience-dependent mating duration of male *Drosophila*": Table 1

**Table 1. Rescue mutant *SIFaR*$^{B322}$ behaviour by using different *SIFaR GAL4* lines.**

| GAL4 | # of Males | # of Females | # of 6B | # of non-6B* |
|---|---|---|---|---|
| 49041 #1 | 18 | 17 | 35 | 0 |
| 49041 #2 | 20 | 15 | 35 | 0 |
| 49061 #2 | 18 | 17 | 19 | 16 |
| 49061 #3 | 13 | 22 | 14 | 21 |
| 49087 #2 | 16 | 17 | 16 | 19 |
| 49087 #3 | 12 | 23 | 17 | 18 |
| 46391 #1 | 148 | 186 | 334 | 0 |

*Presence of non-6B flies is an indicator of rescuing behaviour of lethality phenotype.
