## Supplementary material for "Neuropeptide relay between SIFa signaling controls the experience-dependent mating duration of male *Drosophila*": Table 2

**Table 2. Screening results of neuropeptide GAL4 and RNAi for SIFa signaling**

| NP | ID | GAL4 | RNAi | SIFaR-RNAi (34947) | | SIFaR-GAL4 (49087) | |
|---|---|---|---|---|---|---|---|
| | | | | LMD | SMD | LMD | SMD |
| AKH | Veenstra | GAL4 | | n.s | n.s | | |
| | 34960 | | RNAi | | | ** | n.s |
| | v11352 | | RNAi | | | *** | * |
| AstA | 51979 | GAL4 | | ** | n.s | | |
| | 25866 | | RNAi | | | n.s | n.s |
| AstC | 39448 | GAL4 | | * | n.s | | |
| | 25868 | | RNAi | | | No Data | n.s |
| Burs | 40972 | GAL4 | | n.s | n.s | | |
| | 26719 | | RNAi | | | * | *** |
| | v13520 | | RNAi | | | *** | ** |
| CAPA | 51969 | GAL4 | | n.s | *** | | |
| | 28345 | | RNAi | | | *** | No Data |
| | v41124 | | RNAi | | | n.s | No Data |
| Crz | 51976 | GAL4 | | *** | ** | | |
| | v30670 | | RNAi | | | n.s | n.s |
| DH31 | 51988 | GAL4 | | * | ** | | |
| DH44 | 51987 | GAL4 | | *** | n.s | | |
| | 25804 | | RNAi | | | *** | * |
| | v108473 | | RNAi | | | No Data | *** |
| | 45054 | | RNAi | | | ** | * |
| DSK | 51981 | GAL4 | | ** | *** | | |
| | 25869 | | RNAi | | | n.s | n.s |
| EH | 51974 | GAL4 | | *** | ** | | |
| ETH | 51982 | GAL4 | | *** | n.s | | |
| | 26242 | | RNAi | *** | ** | | |
| FMRFa | 51990 | GAL4 | | n.s | * | | |
| | 58197 | | RNAi | | | n.s | No Data |
| LK | 51993 | GAL4 | | *** | n.s | | |
| | 14091 | | RNAi | | | No Data | n.s |
| MIP | 51984 | GAL4 | | *** | n.s | | |
| | 41680 | | RNAi | | | *** | ** |
| | 5294 | | RNAi | | | *** | ** |
| MS | 51986 | GAL4 | | n.s | * | | |
| | 26245 | | RNAi | | | n.s | n.s |
| Pburs | v27142 | | RNAi | | | n.s | n.s |
| Proc | 51972 | GAL4 | | n.s | n.s | | |
| | 29570 | | RNAi | | | n.s | n.s |
| | v102488 | | RNAi | | | *** | *** |
| sNPF | 46382 | GAL4 | | ** | *** | | |
| TK | 51974 | GAL4 | | ** | * | | |
| | 25800 | | RNAi | | | n.s | n.s |
| | v103662 | | RNAi | | | n.s | *** |
